# Serial dependence transfers between perceptual objects

**DOI:** 10.1101/165399

**Authors:** Greg Huffman, Jay Pratt, Christopher J. Honey

## Abstract

Judgments of the present visual world are affected by what came before. When judgments of visual properties such as orientation are biased in the direction of preceding stimuli, this is called visual serial dependence. Visual serial dependence is thought to arise from mechanisms that support perceptual continuity: because physical properties of an object usually vary smoothly in time, perception might be accurately stabilized by smoothing the perceived features in time. However, mechanisms that support perceptual continuity should be object-specific, because the orientation of one object is more related to its own past than to the past of a distinct object. Thus, we tested the perceptual continuity explanation by comparing the magnitude of serial dependence between objects and within objects. Across three experiments, we manipulated objecthood by varying the color, the location, and both the color and the location of Gabor patches. We observed a serial dependence effect in every experiment but did not observe an effect of objecthood in any experiment. We further observed serial dependence even when the orientations of two successive stimuli were nearly orthogonal. These data are inconsistent with explanations of serial dependence based on visual continuity. We hypothesize that serial dependence arises from a combination of perceptual features and internal response variables, which interact within a common task or decisional context.

---

The physical world changes gradually, and so consecutive states of the world are correlated. For a perceptual system that must infer the origins of sensory input (Clark, 2013; Friston & Kiebel, 2009), it may be adaptive to infer features of the present world to be similar to the recent past. This would lead perceptual report to exhibit a “serial dependence”, a bias to judge present stimuli to be more similar to recent stimuli than they actually were. Serial dependence has been observed for basic perceptual judgments: reports of Gabor patch orientation are biased toward the orientation of stimuli perceived in the previous seconds (Fischer & Whitney, 2014; Fritsche, Mostert, & de Lange, 2017). Serial dependence has also been observed across multiple levels of visual judgment including numerosity (Cicchini, Anobile, & Burr, 2014), facial identity (Liberman, Fischer, & Whitney, 2014), and attractiveness (Taubert & Alais, 2016; Taubert, Van der Burg, & Alais, 2016, Xia, Leib, & Whitney, 2016).

What is the origin and role of these serial dependence phenomena? One hypothesis is that serial dependence phenomena reflect the operation of a “continuity field”, a fundamental mechanism that leads object representation to vary in a continuous manner over time, thus maintaining perceptual stability (Fischer & Whitney, 2014). Here, we test a prediction of the continuity field hypothesis: if serial dependence phenomena arise from a mechanism that maintains perceptual continuity, then serial dependence should be specific to a particular object and its features, and thus should not transfer between distinct objects.

In order to study serial dependence in perception, researchers present series of stimuli that can vary along a continuous dimension and measure the extent to which stimuli from previous trials can influence the perception of the current stimulus. For example, one study had participants judge the orientations of Gabor patch stimuli that differed in orientation trial-by-trial (Fischer & Whitney, 2014). They found that the average orientation judgment for a Gabor patch on a given trial is biased towards to the Gabor patch’s orientation from the previous trial. They found evidence that the bias was perceptual and not response based: no response was needed to the prior Gabor patch for the bias to appear and the bias also appeared when participants completed a simultaneous orientation comparison task. Furthermore, they found that the bias followed attention around a display and dissipated as a function of how far stimuli appeared away from the center of attention. They interpreted these findings as evidence for a serial dependence mechanism that attaches previous perception of items in a given region to the current stimulus, promoting perceptual stability.

The continuity field is defined as the spatial and temporal region in which a given feature is attracted towards that feature of a previous stimulus. The idea of continuity fields as mechanisms for helping maintain perceptual stability has a notable relationship to the notions of feature integration (Treisman & Gelade, 1980) and object files (Kahneman, Treisman, & Gibbs, 1992). Feature integration refers to the idea that, since sensory features are coded in a parallel and distributed fashion (Rogers & Mcclelland, 2014), there must be a mechanism that integrates these independently coded features into individual multi-feature objects. To test for this integrative mechanism, Kahneman et al. presented participants with a “preview display”: an array of placeholder boxes, differentiated by location and/or color, with a letter in each of them. The letters then offset before the boxes moved to new locations. Next, a single letter appeared within one of the boxes. This newly-appeared letter could be the same letter that appeared in that box during the preview, the same as a letter that appeared in a different box during the preview, or a letter that was absent from the preview display. Kahneman and colleagues hypothesized that if features are bound into specific object files there should be a larger benefit for identifying letters appearing within the same box twice (object specific preview effect) compared to responding to letters that had appeared in both displays, but in different placeholders (object non-specific preview effect), or letters absent from the preview display. The data supported this hypothesis across a range of conditions. The authors concluded that there is a feature integration mechanism that binds independent features into object files, leading to a cost when an unexpected letter appears in a placeholder.

How does the serial dependence phenomenon relate to object-specific preview effects? In both cases, past perception influences current perception, and in both cases these effects depend on the similarity of past and present stimuli. One difference between the phenomena, as noted by Fischer & Whitney (2014), is that repeating objects leads to more accurate object identification on the second presentation, while serial dependence studies show that current perception is made less accurate by previous experience. This difference is not conclusive, of course, because the object identification studies were based on categorical judgments about the second object, rather than measurements of features along continuous dimensions. Still, Fischer and Whitney tentatively suggested that the continuity field (used to explain serial dependence phenomena) may provide a basic mechanism necessary for the creation and updating of object files (used to explain object tracking phenomena).

If a continuity field introduces a bias to group and track object features, then it should not bias features of one single object in the same way as two different objects. Features of a single object are expected to be auto-correlated in time, and so perception may be usefully biased by assuming continuity. In contrast, features of distinct objects may be completely uncorrelated in time, and assuming continuity of features across objects can lead to erroneous perception. Thus, we can test the continuity field explanation by measuring the magnitude of serial dependence while manipulating factors that cause stimuli to be treated as separate objects. If continuity field mechanisms drive serial dependence, then these effects should be limited to individual objects, or at least stronger within objects than across objects. However, if serial dependence is not affected by object membership, then it seems implausible that it arises from a perceptual continuity field or from other mechanisms that support object files or object tracking.

In order to manipulate objecthood in the current study we manipulated stimulus location and color. We chose to manipulate location and color because they modulate object perception in wide-ranging visual performance tasks. For example, Hommel (1998) had participants make a precued left or right response to a red or green X or O stimulus appearing above or below fixation. Next, the participants needed to identify a second red or green X or O using the same response keys. He found that location and color repetitions between the first and second stimulus affected object identification, even when they were task irrelevant. Color and/or location play a role in object perception even when minimally attended to (Hommel, 2005), when objects are presented subliminally (Keizer, Hommel, & Lamme, 2015), and when more complex stimuli are used (Golomb, Kupitz, & Thiemann, 2014). Furthermore, location is automatically integrated with other stimulus features (Hommel, 2002), may be preferentially processed (Chen, 2009) and may determine which non-spatial features are integrated together (Kahneman, Treisman, & Gibbs, 1992; Treisman & Gelade, 1980; van Dam & Hommel, 2010). Indeed, even when color is outside of an individual’s attentional control setting (Folk, Remington, & Johnston, 1992) it can lead to object-based processing effects (Carmel & Lamy, 2014; Huffman, Antinucci, & Pratt, 2018; Schoeberl, Ditye, & Ansorge, 2018). As a whole, within the feature integration literature, stimulus location and color have been found to be powerful features for differentiating objects.

To test how serial dependence varies with object membership, we employed a variation of the Fischer and Whitney (2014) procedure in which stimulus features were manipulated so that sequentially presented stimuli were understood as being part of the same or different objects. In Experiment 1, we presented the Gabor patches in blue or yellow, reasoning that individuals would treat differently colored stimuli as different objects. In Experiment 2, we presented the target stimuli to the left or right of fixation randomly on each trial, reasoning that individuals would treat consecutive stimuli at separate locations as separate objects. Finally, in Experiment 3, we manipulated both stimulus color and location such that one color stimulus would always appear at one location and the second stimulus would always appear at the other location. In all cases, we asked participants to report the Gabor patch’s orientation and measured the extent to which any error was biased towards the orientation of the previous stimulus. To anticipate our findings: we found the serial dependence effect, but did not find any effect of whether the previous stimulus was likely perceived as the same object or a different object.

## Experiment 1: Manipulating Stimulus Color

In Experiment 1, a Gabor patch always appeared at the same location (within a block of trials) but its color could switch between consecutive trials. We asked: does the orientation of a stimulus on a given trial affect the orientation judgments of the next stimulus only when they are the same color (and might be the same object) or does it also transfer when the stimuli are two different colors (and are unlikely to be the same object)? If the serial dependence effect is sensitive to objecthood, we predict a significantly reduced serial dependence effect when stimulus color switches across trials (Figure 1).

**Figure 1.**
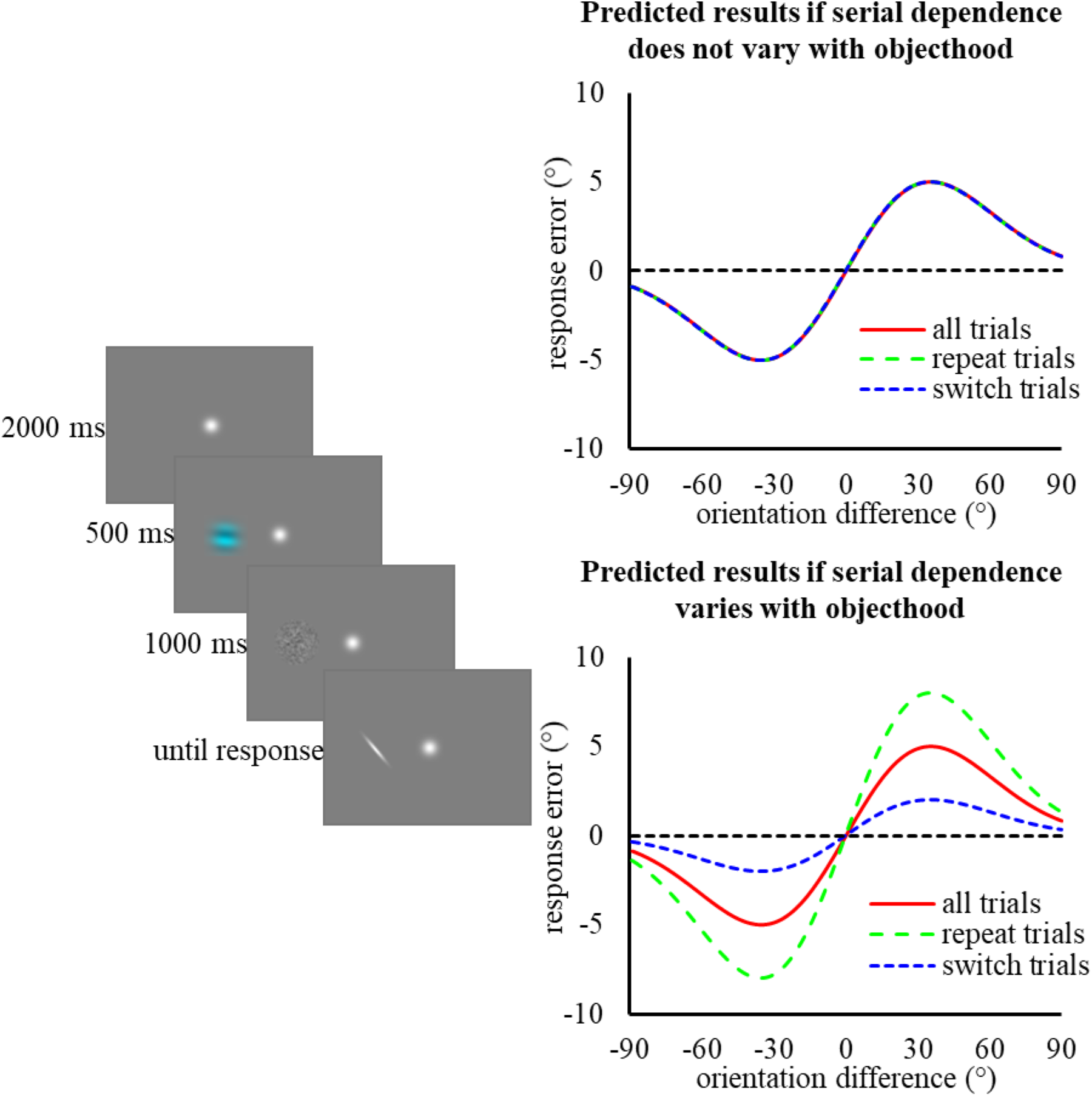
Left panel: Stimuli and procedure. Participants first viewed a Gabor patch to the left or right of fixation (images are not to scale). This was followed by a white noise mask. Participants then saw a response bar which they rotated to the same orientation as the Gabor patch, using the arrow keys. Right panel: Predicted results if serial dependence does not (top panel) or does (bottom panel) vary as a function of objecthood.

## Methods

### Participants

Seven undergraduate students from the University of Toronto participated in the experiment and we compensated them $10 per hour of participation. All participants provided informed consent prior to the experiment and reported normal or corrected-to-normal visual acuity and normal color vision. Experiments were done in agreement with the Declaration of Helsinki.

### Stimuli and apparatus

Participants completed the experiment using a PC connected a CRT monitor (screen resolution: 1024 × 768; refresh rate: 85 Hz). We used MATLAB (Mathworks) with the Psychophysics toolbox (Kleiner, Brainard, Pelli, Ingling, Murray, & Broussard, 2007) for stimulus presentation. Stimuli appeared on a gray background. The target Gabor patch stimuli were sine wave gratings with a spatial frequency of 0.33 cycles per degree presented in a 1.5° *SD* Gaussian contrast envelope. We used white noise patches to decrease aftereffects. The white noise patches were smoothed with a 0.20° Gaussian kernel and windowed in a 1.5° *SD* Gaussian contrast envelope. The Gabor patches could be blue (RGB: 000, 000, 255) or yellow (RGB: 255, 255, 000). The fixation target was a white dot windowed in a 0.5° *SD* Gaussian contrast envelope. The response bar stimulus was a white 0.61° white bar windowed in a 1.5° *SD* Gaussian contrast envelope. The Gabor patch stimuli were presented 6° to the left or right of fixation (manipulated between blocks). Participants responded using the left and right arrow keys on a QWERTY keyboard. We presented the response bar at a random orientation. We had participants use chin and head rests to maintain an approximate viewing distance of 44 cm.

### Procedure

We presented the fixation target at the center of the display, where it remained throughout the experiment. We instructed participants to maintain fixation throughout the experiment. After 2000 ms, a Gabor patch of random orientation was presented either 6° to the left or to the right of the fixation stimulus, on the horizontal midline of the display. The Gabor patch remained for 500 ms before offsetting and was followed by the white noise stimulus that remained for 1000 ms. After 250 ms, the response bar appeared at the same location as the Gabor patch stimulus. Using the left and right arrow keys, participants then rotated the response bar to match the orientation at which they perceived the Gabor patch stimulus. Once they were satisfied with the response bar’s orientation, they pressed the spacebar to end the trial.

### Design

Participants completed eight blocks of 140 trials (70 yellow Gabor patch trials, 70 blue Gabor patch colors) over the course of three sessions on different days for a total of 1120 trials. Across trial blocks the Gabor patch alternated between appearing to the left or right of fixation. Within a block the Gabor always appeared at the same location. We did not counterbalance the number of color repeats/switches. The orientations of the Gabor patch and the response bar were determined randomly on each trial: the orientations were sampled independently from a uniform distribution on the range [0, 180) degrees.

### Analyses

For each trial, *n*, of the experiment we defined two categorical predictor variables: one predictor tracked whether the color switched from trial *n*-1 to trial *n*; the other predictor reflected the difference in orientation between the stimulus on trial *n* and the stimulus on trial *n*-1. The orientation difference (orientation on trial *n* minus orientation on trial *n-1*) was binned into one of 12 bins, each of 15° width, with bin centers at: −75°, −60°, −45°, −30°, −15°, 0°, 15°, 30°, 45°, 60°, 75. The dependent variable on trial *n* was the “response error”, which was the veridical presented orientation on trial *n* minus the subject-reported orientation on trial *n*.

If some form of serial dependence was present in the data, then the response error on trial *n* should depend on the stimulus that was presented on trial *n-1*. Thus, to test for serial dependence, we tested whether the response error was the same across all orientation difference bins. To instantiate the null hypothesis that the distributions of response errors have the same mean across all bins, we used a permutation test. For each bin, we (i) generated surrogate data for our bin by randomly permuting across trials the orientation difference value assigned to each trial and (ii) calculated the mean response error for this surrogate data. This surrogate sampling procedure was repeated 10,000 times to produce a distribution of surrogate values of the mean response error, under the null hypothesis that response errors do not vary across bins. If fewer than 5% of the surrogate mean errors in the null distribution were greater than the observed mean error, we took this as evidence that the response errors in that bin were different from the mean response errors across bins. This statistical procedure was performed for every orientation difference bin, using data pooled from color repeat and color switch trials. The analysis was then performed for each orientation bin, separately for color repeat trials and for color switch trials.

To measure whether the serial dependence differed for repeat and switch trials in each bin, we again used a permutation procedure. We first combined the data from negative and positive orientation difference bins, switching the sign of the response error from negative orientation bin trials (since the serial dependence effect is symmetrical). We then generated repeat-switch trial differences under the null hypothesis by (i) shuffling the condition labels (switch/repeat) across the entire data set (across subjects) and recorded the difference in the serial dependence effect for repeat trials and switch trials. Repeating this permutation procedure 10,000 times we generated a null distribution of repeat-switch difference values. We considered there to be a significant effect of objecthood if the observed repeat-switch difference within a given orientation bin was outside the inner 95% of values obtained through the shuffling procedure.

If the serial dependence effect is reduced by switching color, this should manifest as an overall reduction in the magnitude of response error. Under the serial dependence model, the response error is different for clockwise and counter-clockwise orientation differences. Therefore, if the serial dependence effect is reduced by switching color, this reduction in response error should occur in different directions for clockwise orientation difference trials and for counter-clockwise trials. Furthermore, we expected that the switch-repeat differences should be clearest at the orientations where serial dependence effects are largest: for orientation differences between 15 and 45 degrees (Fischer & Whitney, 2014). Therefore, to directly test this phenomenon at the single-subject level: (i) within each subject, we extracted all data from the - 15, −30, and −45 orientation difference bins (clockwise bins) bins and all data from the 15, 30, and 45 orientation difference bins (counter-clockwise bins); (ii) for both clockwise and counter-clockwise bins separately, the response error on the *n*-th presented color repeat trial was paired with the *n*-th presented color switch trial within a block; (iii) we computed the difference between all repeat-switch pairs; (iv) separately for the clockwise and counter-clockwise data, we computed 95% confidence intervals on the mean of the switch effects. Confidence intervals on the mean of each distribution were computed by generating a distribution of the means across 10,000 surrogate datasets, each generated by sampling with replacement from the empirical distribution. This process of computing means and confidence intervals on the switch effects was performed separately for each subject and separately for the clockwise and counter-clockwise bins within each subject.

Finally, following the analytical method used by previous serial dependence studies (e.g. Fischer & Whitney, 2014) we modelled the data with a derivative-of-Gaussian curve, *y*(*x*) = *awxce*^−(*wx*)2^. We fit the repeat trials and switch trials separately using the fminsearch function in MATLAB minimizing the sum of squared error. If the serial dependence effect is larger in repeat trials, then the amplitude of the curve fit (parameter *a*) to the repeat data should be larger than the amplitude of the curve fit to the switch data. We also tested for whether the serial dependence effect differed between the repeat and switch conditions within individual participants. To do so, we created surrogate data sets by scrambling the repeat/switch condition labels (within each participant), we fit derivative-of-Gaussian curves to the repeat and switch data, and we recorded the difference in amplitude between the two conditions. This was repeated 10,000 times. If the observed repeat-switch difference for a given participant was larger than 95% of the difference found in the permutated data, then the participant showed significantly different serial dependence effects in the repeat and switch conditions.

## Results

Overall, the mean response error was small (*M* = 0.83°) with small standard deviation (*SD* = 14.8°) reflecting accurate performance. As can be seen in the left panel of Figure 1, we observed an overall serial dependence effect. When stimuli on consecutive trials were nearly the same orientation, we observed little response error. As stimulus orientation became increasingly different from the previous stimulus orientation the response error became significantly biased in the direction of the previous stimulus orientation, consistent with the serial dependence effect. In particular, the observed response error was significantly different from zero in the −75° (*M* = - 1.89, *p* < .0001), −60° (*M* = −3.21, *p* < .0001), −45° (*M* = −1.87, *p* < .0001), −30° (*M* = −3.85, *p* < .0001), 15° (*M* = 3.52, *p* < .0001), 30° (*M* = 4.24, *p* < .0001), 45° (*M* = 3.03, *p* = .0007), and 75° (*M* = 3.73, *p* < .0001) bins. The observed response error was not different from zero in the −15° (*M* = −0.169, *p* = 0.0728), 0° (*M* = −0.0154, *p* = 0.106), and 60° (*M* = −1.75, *p* = 0.910) bins. Notably, inconsistent with previous serial dependence research, the mean response error did not return to baseline at either the −75° or 75° orientation bins.

Of most relevance to our central research question is the result that switching object color did not reduce the serial dependence effect. We observed no systematic differences due to color repeat/switch (Figure 2). The mean of repeat response error minus switch response error was indistinguishable from zero (Bonferroni corrected α = .0083) in the 0° (*M* = 1.38°, *p* = .111), 15°, (*M* = 0.445, *p* = .608), 30° (*M* = 0.209, *p* = .807), 45° (*M* = 0.794, *p* = .359), 60° (*M* = 0.291, *p* = .734), and 75° (*M* = −0.759, *p* = .381) bins.

**Figure 2.**
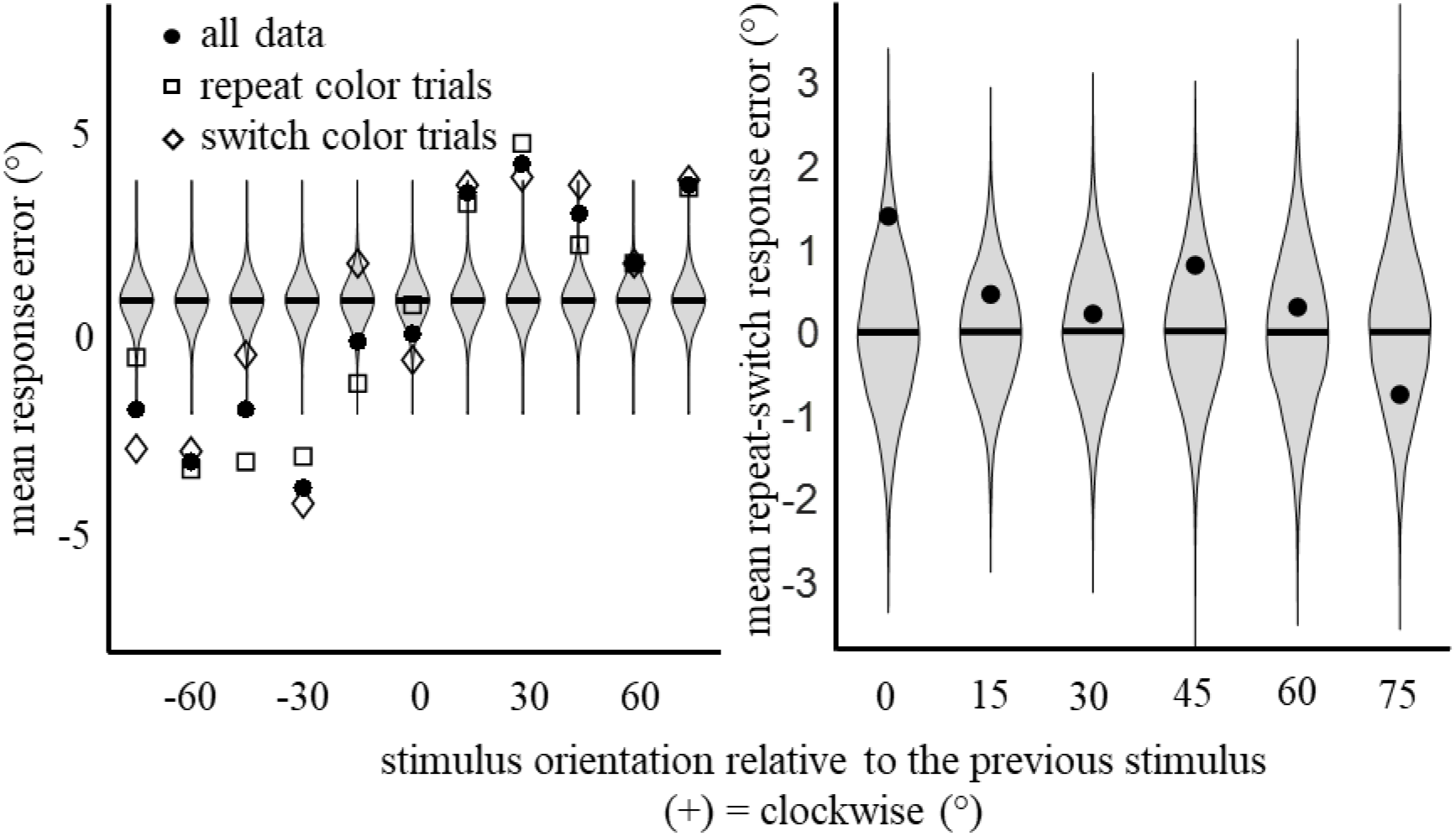
Serial dependence is not detectibly affected by color switches. Left panel: The mean response error as a function of orientation difference for Experiment 1. Violin plots show distributions of response error under the null hypothesis that error is independent of the previous trial’s orientation. Right panel: The mean response errors in color repeat trials minus the mean response error on color switch trials. The violin plots show the distribution of response error under the null hypothesis that response error is independent of whether objecthood repeated or switched.

That the serial dependence effect was unaffected by color repetition was further supported by the derivative-of-Gaussian analysis. The mean difference between the amplitudes of the curves fit to the repeat and switch data was 0.144° (*SE* = 0.715°), which did not differ from zero (paired samples *t*-test comparing the subjects amplitude estimates in the repeat and switch conditions), *t*(6) = .201, *p* = .847. Furthermore, no individual participants demonstrated a significant different serial dependence effect based on color repetition condition (all *p*s > 0.114). Using JASP (JASP Team, 2018), we conducted a Bayesian, one sample *t*-test with a Cauchy prior and with the scale setting of 1. This test indicated that the null hypothesis was 3.63 times more likely than the alternative hypothesis that objecthood affects the serial dependence effect.

Finally, we checked in individual subjects for an effect of color switching, focusing on the 15°-45° range of inter-trial shifts, for which serial dependence effects are largest. This analysis revealed no significant differences between repeat and switch trials for the clockwise or counterclockwise rotation bins for any participants (Figure 3). For all subjects, and for both the positive and the negative orientation bins, the 95% confidence intervals on the serial dependence effect overlapped for switch and repeat bins.

**Figure 3.**
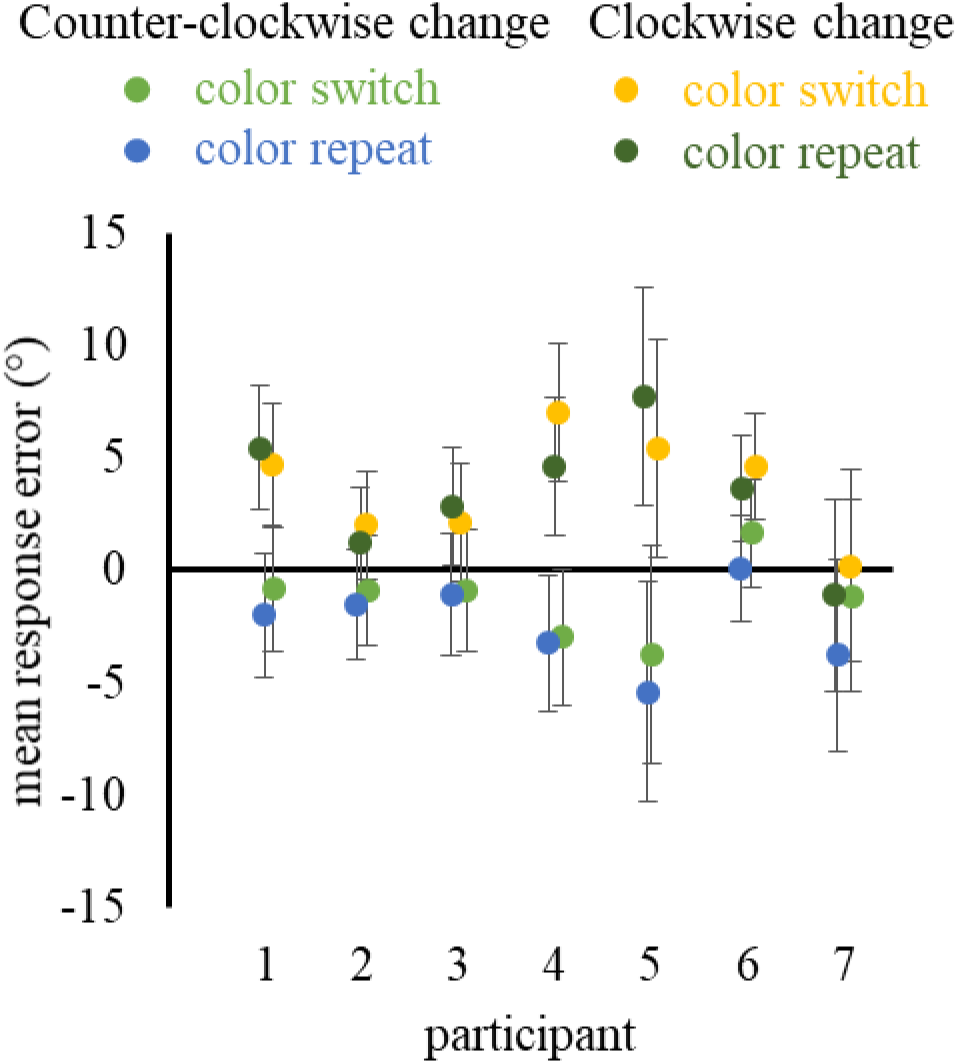
The mean response error aggregated across the +/- 15° - 45° orientation bins, separated for individual subjects and for color repeat and color switch trials. The error bars represent the 95% confidence intervals of the repeat – switch response error difference (generated through resampling).

## Discussion

The data from Experiment 1 revealed a serial dependence effect: orientation judgments were systematically biased toward the orientation of the previous stimulus. However, this serial dependence effect was not detectibly affected by whether the stimulus repeated or switched colors between trials. This finding is inconsistent with the notion that serial dependence reflects a mechanism for object-level perceptual continuity. A second, incidental finding, was that the response errors were significantly different from zero even at the ±75° orientation difference bins. This observation stands in contrast to both Fischer and Whitney (2014) and Fritsche and colleagues (2017) who found that the response error returned to zero or turned negative, in some cases. This may indicate that the interleaved color change manipulation is somehow affecting the serial dependence effect. It is unclear why more difference between stimuli would lead the serial dependence mechanism to apply across even more dissimilar stimuli.

## Experiment 2: Manipulating Stimulus Location

Although we observed a serial dependence effect in Experiment 1, we also found that it existed even at the most extreme orientation bins, a result that differs from Fischer & Whitney (2014). Therefore, in Experiment 2 we used location rather than color to manipulate objecthood. By removing the color switch, we more closely approximate the Fischer and Whitney (2014) design, and are once again to test for an effect of objecthood. In particular, we used grayscale stimuli that could appear at either side of fixation. Since perceptual objects are addressed by their location (Kahneman, Treisman, & Gibbs, 1992), this manipulation may be more likely to lead to repeat – switch differences. Additionally, when the stimuli repeat at the same location, the design is essentially the same as the first experiment by Fischer and Whitney. This allows us to test whether that response errors continue to be biased at the ±75° orientation bins even while stimuli repeat in color and location as in prior studies.

## Methods

### Participants

Six undergraduate students from the University of Toronto participated in the experiment and we compensated them $10 per hour of participation. All participants provided informed consent prior to the experiment and reported normal or corrected-to-normal visual acuity and normal color vision.

### Stimuli and apparatus

The stimuli and apparatus replicated those from Experiment 1, except that all Gabor patches were presented in grey.

### Procedure and design

The procedure matched the procedure of Experiment 1 with the following exceptions. The stimuli were greyscale and could appear 7° to the left or right of fixation. Within each trial block of 140 trials the stimulus appeared on the left for 70 trials and the right for 70 trials and participants completed nine trial blocks rather than eight. We did not counterbalance the number of location switches/repeat, so there may have been small random differences across subjects in the number of repeat/switch trials occurring on the left/right.

## Results

Overall, the mean response error was small (*M* = 1.36°) with small standard deviation (*SD* = 11.6°) reflecting accurate performance. Once again, we successfully replicated the essential features of the serial dependence effect (Figure 4). When stimuli on consecutive trials were nearly the same orientation, we observed little response error. Specifically, the observed response error was significantly different from zero in the −75° (*M* = −0.930, *p* < .001), −60° (*M* = −1.46, *p* < .001), −45° (*M* = −0.421, *p* < .001), −30° (*M* = −2.81, *p* < .001), −15° (*M* = −1.87, *p* < .001), 15° (*M* = 4.74, *p* < .001), 30° (*M* = 3.77, *p* < .001), 45° (*M* = 3.08, *p* = .0010), and 75° (*M* = 2.94, *p* = .0014) bins. The observed response error was not different from zero in the 0° (*M* = 0.827, *p* = .158) and 60° (*M* = 1.57, *p* = 0.646). As stimulus orientation became increasingly different from the previous stimulus orientation the response error became significantly biased in the direction of the previous stimulus orientation. As in Experiment 1, the response error in the repeat condition did not return to the baseline mean response error in the −75° orientation bin or the 75° orientation bin it approached the baseline error

**Figure 4.**
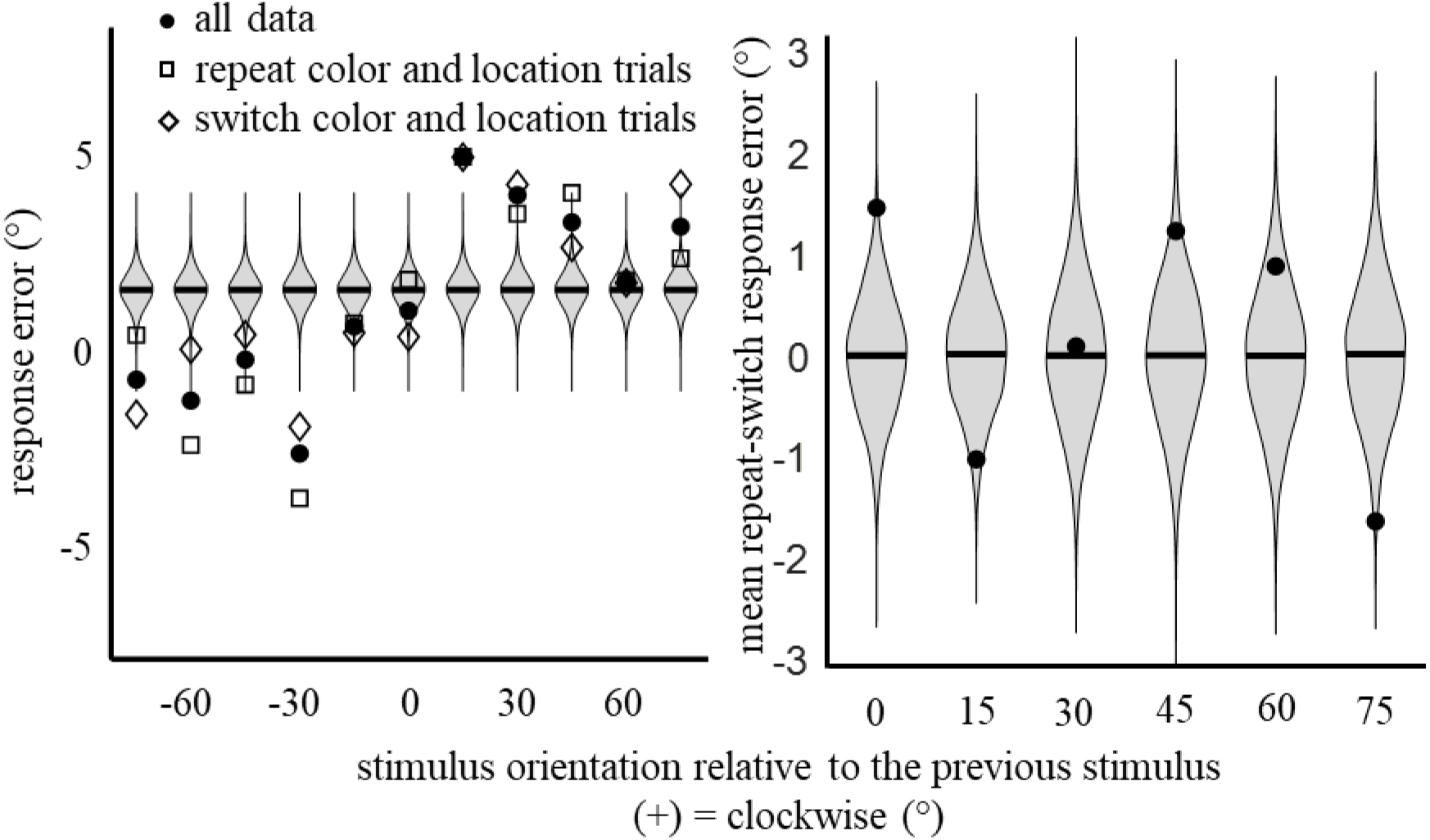
Serial dependence is not detectibly affected by color switches. Left panel: The mean response error as a function of orientation difference for Experiment 1. Violin plots show distributions of response error under the null hypothesis that error is independent of the previous trial’s orientation. Right panel: The mean response errors in color repeat trials minus the mean response error on color switch trials. The violin plots show the distribution of response error under the null hypothesis that response error is independent of whether objecthood repeated or switched.

In terms of the question as to whether objecthood affects the magnitude of serial dependence, we did not observe any systematic differences due to the location switch (Figures 4). The mean repeat – switch response error was not different from zero (Bonferroni corrected α = .0083) in the 0° (*M* = 1.45°, *p* = 0.0138), 15° (*M* = −1.03°, *p* = 0.0837), 30° (*M* = 0.0874°, *p* = 0.885), 45° (*M* = 1.23°, *p* = 0.0404), and 60° (*M* = 0.878°, *p* = 0.136) orientation bins. There was a significantly larger serial dependence effect in the repeat compared to switch condition in the switch compared repeat condition in the 75° (*M* = −1.64°, *p* = 0.0063) orientation bin.

We also measured at the repeat – switch difference within individual participants in the clockwise and counterclockwise rotation bins separately (using only the 15° - 45° bins within each). As can be seen in Figure 5, response errors in the positive and negative orientation bins were indistinguishable in the repeat and switch conditions for all subjects.

**Figure 5.**
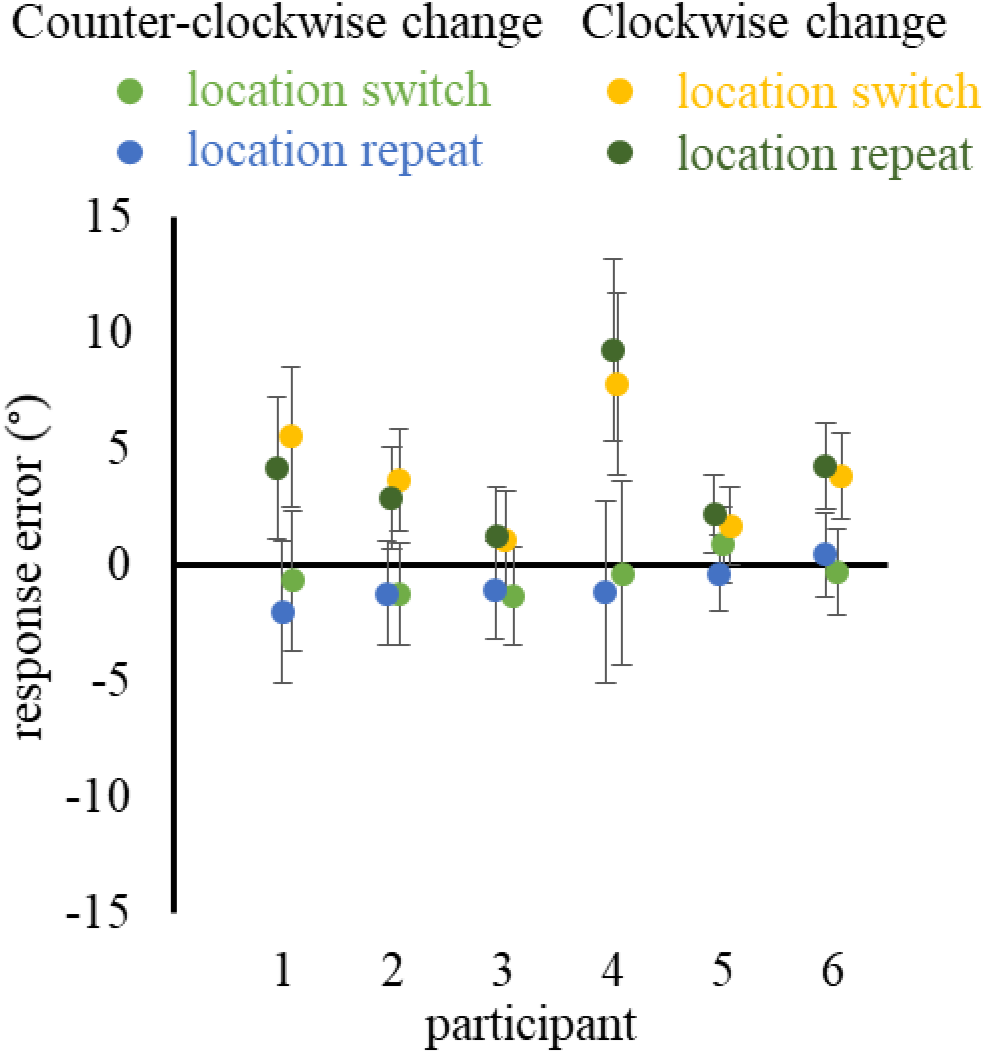
The mean response error aggregated across the +/- 15° - 45° orientation bins, separated for individual subjects and for location repeat and location switch trials. The error bars represent the 95% confidence intervals of the repeat – switch response error difference. Confidence intervals were generated by resampling.

Finally, the derivative-of-Gaussian analysis found no difference in the serial dependence effect when location was repeated versus. switched. The mean location repetition effect was - 0.64° (*SE* = 0.29°), which did not differ from zero, *t*(5) = 2.228, *p* = .076. At the individual participant level, one participant showed a 4.13° larger serial dependence effect in the repeat condition, *p* = .0270. Another participant showed a 1.71° larger serial dependence effect in the switch condition, *p* = .0249. A Bayesian one-sample *t*-test indicated that the alternative hypothesis was 1.434 times more likely, but in the opposite of the predicted direction.

## Discussion

As in Experiment 1, we observed a systematic serial dependence effect. We further found that the serial dependence effect was not detectibly different when the stimulus location repeated or switched across trials. In Experiment 1 we found larger effects in the switch condition in the - 45° and −15° orientation bins; in Experiment 2 we found that difference for the 75° bin, but nowhere else. The fact that these differences are small, and of inconsistent direction, combined with the absence of differences for any individual subject (Figure 5), suggest that there is no systematic effect of switching location. Given this, our next step was to test whether an even more salient manipulation of objecthood would reduce the serial dependence effect.

It is perhaps unsurprising that manipulating location did not affect the serial dependence effect, as Fischer and Whitney (2014) found that the effect did not dissipate until stimulus location differed by more than 20°. That said, the stimuli in that previous experiment appeared at random locations across the visual display. It is possible that participants needing to monitor the entire display for stimulus onsets may lead to the continuity field being more broadly tuned than it would be in other contexts. In contrast, by using two locations we more closely replicate studies that did find location-based object file effects (Colzato & Hommel, 2004; Hilchey, Rajsic, Huffman, Klein, & Pratt, 2018; Hommel, 1998). The absence of a location switch/repeat effect in this experiment demonstrates the location-invariance of the serial dependence effect, under conditions similar to those in which location switch/repeat is thought to affect the access of object files.

## Experiment 3: Manipulating Stimulus Color and Location

In Experiment 1 we manipulated objecthood by changing the color of the stimulus across trials whereas in Experiment 2 we did so my manipulating stimulus location. Neither manipulation altered the serial dependence effect. In Experiment 3, we manipulate both these factors simultaneously in order to maximize the differences between the two objects and increase the possibility of objecthood affecting the serial dependence effect. In this experiment, either a blue or a yellow stimulus was presented on each trial, but a blue stimulus would always appear left of fixation and a yellow stimulus would appear right of fixation. Given the suggestion that objects are “addressed” by their locations (Kahneman, Treisman, & Gibbs, 1992) and color is typically a salient feature (Theeuwes, 1992), this procedure may lead to stronger object differentiation and should prevent the serial dependence effect from transferring between objects.

## Methods

### Participants

Seven undergraduate students from the University of Toronto participated in the experiment and we compensated them $10 per hour of participation. All participants provided informed consent prior to the experiment and reported normal or corrected-to-normal visual acuity and normal color vision.

### Stimuli and apparatus

The stimuli and apparatus replicated those from Experiment 1.

### Procedure and design

The procedure matched that of Experiment 1, except for the location of the stimulus. Rather than blue and yellow Gabor patches appearing at the same location on each trial, each color always appeared on one side of the fixation stimulus or the other. That is, for example, the blue stimulus always appeared on the left and the yellow stimulus always appeared on the right. Whether the left/right stimulus was blue/yellow was alternated across trial blocks. We did not counterbalance the number of location/color repeats/switches. Observers completed nine blocks of 140 trials, leading to a total of 1260 trials per observer.

## Results

Overall, the mean response error was small (*M* = 0.11°) with small standard deviation (*SD* = 11.8°) reflecting accurate performance. As can be seen in the left panel of Figure 6, we again found a serial dependence effect. Specifically, the observed response error was significantly different from zero in the −75° (*M* = −0.948°, *p* = 0.0094), −60° (*M* = −1.47°, *p* = .0001), −45° (*M* = −1.95°, *p* < .0001), −30° (*M* = −2.51°, *p* < 0.0001), 15° (*M* = 4.03°, *p* < .0001), 30° (*M* = 2.06°, *p* < .0001), 45° (*M* = 2.83°, *p* < .0001), 60° (*M* = 1.28°, *p* = .0031), 75° (*M* = 1.43°, *p* = .0012). Once again, the overall response error in the ±75° orientation bins did not return to baseline.

As for the repeat – switch difference, Figure 6 (right panel) illustrates that the repeat – switch response error was not different from zero (Bonferroni corrected α = .0083) in the 0° (*M* = −0.443°, *p* = 0.440), 15° (*M* = 0.123°, *p* = 0.826), 30° (*M* = −0.929°, *p* = 0.107), 45° (*M* = −1.124°, *p* = 0.0536), 60° (*M* = −1.409°, *p* = 0.0142), and 75° (*M* = −0.183°, *p* = 0.746) orientation bins.

**Figure 6.**
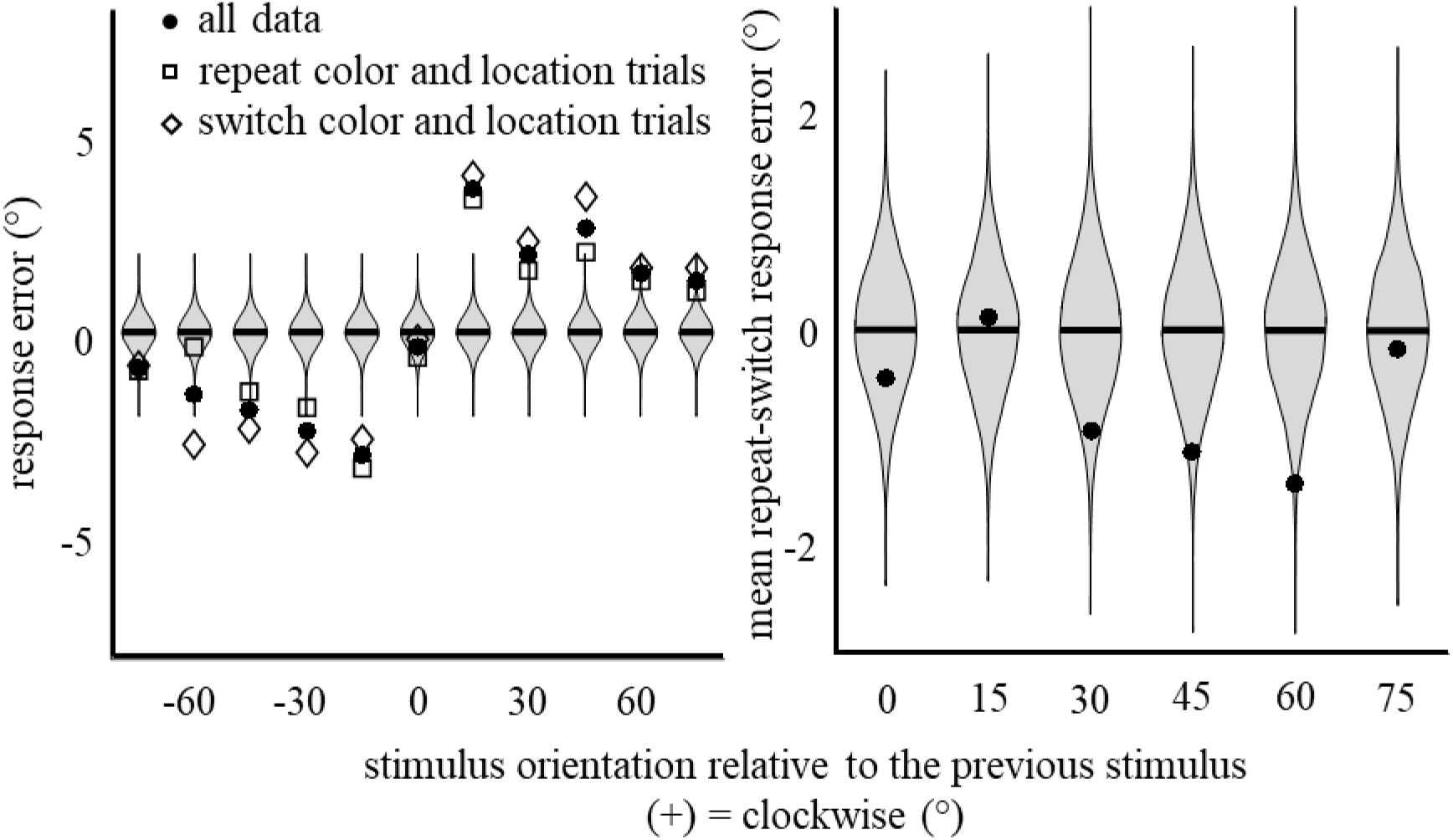
Serial dependence is not detectibly affected by color/location switches. Left panel: The mean response error as a function of orientation difference for Experiment 1. Violin plots show distributions of response error under the null hypothesis that error is independent of the previous trial’s orientation. Right panel: The mean response errors in color repeat trials minus the mean response error on color switch trials. The violin plots show the distribution of response error under the null hypothesis that response error is independent of whether objecthood repeated or switched.

We again examined the repeat – switch difference within individual participants in the clockwise and counterclockwise rotation bins separately. As can be seen in Figure 7, this analysis revealed no significant differences between repeat and switch trials for the clockwise or counterclockwise rotation bins for any participants.

**Figure 7.**
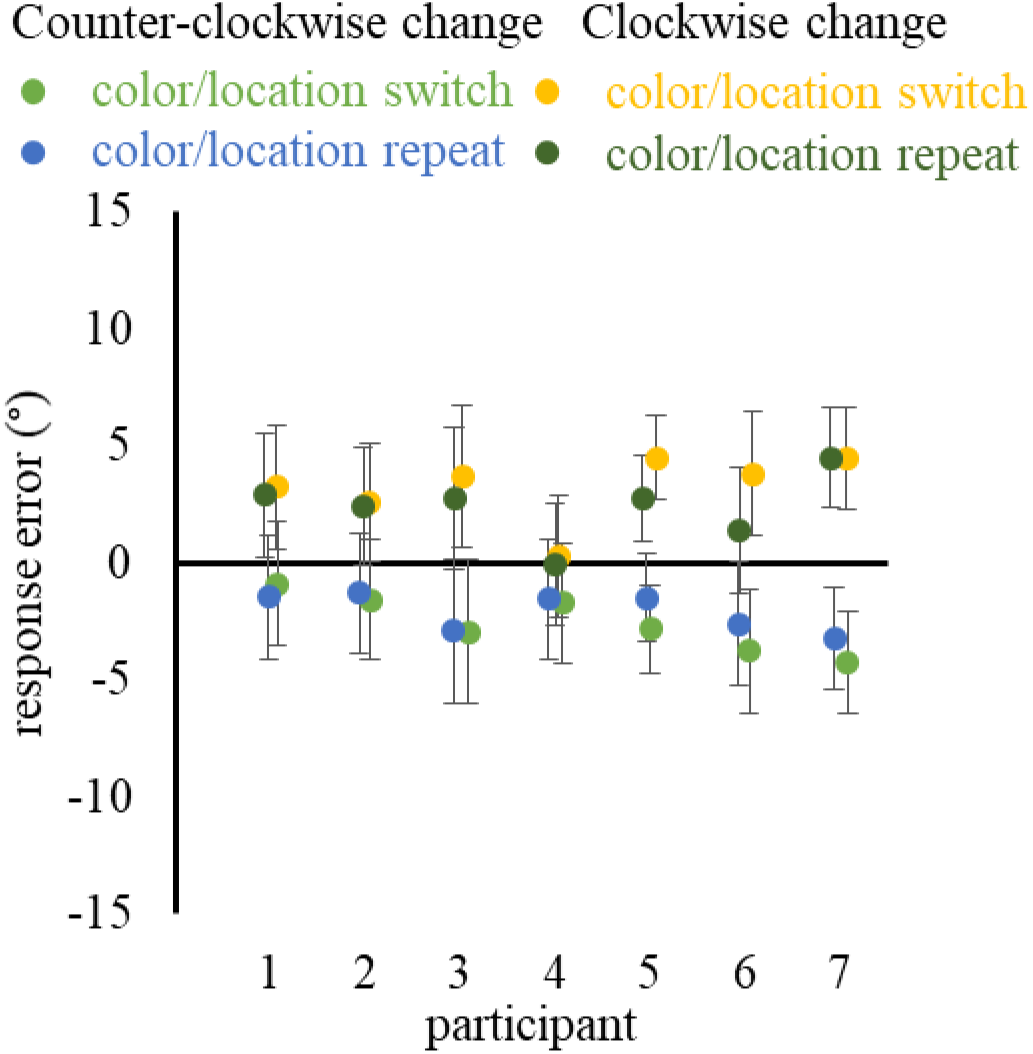
The mean response error broken done by subject and color repeat/switch using only the +/- 15° - 45° orientation bins. The error bars represent the 95% confidence intervals of the repeat – switch response error difference (generated through resampling).

Finally, the derivative-of-Gaussian analysis again revealed no evidence that the serial dependence effect differed by color/location repetition. The difference between the color/location repeat and switch conditions was 0.37° *(SE* = 0.75°), which did not differ from zero, *t*(6) = 0.492, *p* = .640. No individual participants showed a different serial dependence effect based on the color/location repetition condition, *p* > 0.15. A Bayesian, one-sample t-test indicated that the null hypothesis was 2.56 times more likely than the alternative hypothesis that objecthood affects the serial dependence effect.

## Discussion

The results of Experiment 3 were consistent with those of Experiments 1 and 2. In this case, we observed a serial dependence effect that was object non-specific even when consecutive stimuli were differentiated by a change in color and a change in location. In addition, the mean response errors remained significantly different from zero even at the ±75° orientation difference bins.

## General Discussion

Previous studies have observed serial dependence in perception. These dependences are hypothesized to result from continuity fields: regions of space in which perceived stimulus features are biased in the direction of previous stimuli within that space. In this way, the continuity field can support perceptual continuity over time. To be adaptive for visual perception, however, the biasing effect should be applied only to stimuli that are sufficiently similar to previous stimuli within the field. If one object’s features ubiquitously biased the features of a separate object, then the continuity field would become maladaptive. Therefore, we tested whether the continuity field was invariant to object-level similarity by manipulating factors that cause stimuli to be perceived as different perceptual objects.

Our study produced two consistent findings. First: we observed no reliable reduction in serial dependence between stimulus pairs that should be perceived as different objects. We manipulated color (Experiment 1), location (Experiment 2), and color and location simultaneously (Experiment 3). For each manipulation we observed that serial dependence was indistinguishable for consecutive stimuli that should be perceived as the same object or as different objects. Second, we found evidence the serial dependence effect arises from an interaction between perceptual features and internal response variables: responses were weakly biased towards the previous stimulus even when that previous stimulus had a very dissimilar orientation (more than 70 degrees rotated) from the current stimulus. This was the case in all three experiments, including the repeat condition of Experiment 2 which closely approximates Experiment 1 from Fischer and Whitney (2014).

Our interpretation of these results rests on the assumption that manipulating color and location is sufficient for causing individuals to perceive two stimuli as two different objects. We chose to manipulate color and location because of the extensive evidence that these two features are integrated into object representations, even when task-irrelevant (e.g., Hommel, 1998; Hommel & Colzato, 2004; Hommel, Memelink, Zmigrod, & Colzato, 2013; Quinlan, 1998).

This is the case in stimulus identification tasks (Hilchey, Rajsic, Huffman, & Pratt, 2017a, 2017b), in priming of pop-out tasks (Krummenacher, Müller, Zehetleitner, & Geyer, 2009), and in attentional cueing tasks (Huffman, Al-Aidroos, & Pratt, 2016; Huffman, Antinucci, & Pratt, 2018). The effects of location and color persist even when manipulations are applied in the attempt to eliminate object-based effects (Hilchey, Rajsic, Huffman, & Pratt, 2017a, 2017b). Overall, the strength of object-integration effects relating to color and location across diverse paradigms support our assumption that objects appearing in different locations and/or colors were perceived as different objects. Beyond our manipulation of the target’s visual features, it is notable that two targets were separated by more than 3000 ms and during that interval a white noise mask and adjustment bar appeared at the target’s location. So, the targets were temporally segregated and the stimulus last associated with a target’s location was the response bar, not the target stimulus.

While our findings are difficult to reconcile with a purely perceptual model for the serial dependence phenomenon, they are consistent with models that combines perceptual and decisional variables. The original explanation for serial dependence emphasized perceptual contributions, via the notion of a “continuity field” (Fischer & Whitney, 2014): a region of space in which the visual system assumes that stimuli close in space and time have similar visual properties (e.g., that two consecutive Gabor patches should have similar orientation). Assuming continuity has the benefit of enhancing continuity (smoothness of change) in perception. Also, under noisy conditions, the continuity assumption can enhance the accuracy of perception, as long as the continuity assumption is correct. However, the continuity assumption is less likely to be correct when consecutive visual properties are sampled from distinct objects: physical properties of the world vary gradually over time within objects (e.g., a rotating soccer ball), but can vary discontinuously across distinct objects (e.g., two different soccer balls). Substantial evidence indicates that the continuity field must arise from a perceptual contribution and is not a purely decisional (or even motoric) phenomenon. A purely motoric account can be ruled out by the fact that serial dependence persists even when a motor response is not required. Fischer & Whitney (2014) interleaved trials that did and did not require a motor response. They found that serial dependence was equally large when no motor response was produced for the preceding stimulus.

What is the evidence for a perceptual contribution to serial dependence? Firstly, the magnitude of serial dependence depends on the similarity of the present and prior visual features, a factor that is independent to the response decision on each trial. In the present experiments, we did not observe serial dependence returning to baseline for near-orthogonal stimuli, but we did replicate the finding that that serial dependence was strongest effects for 15-45° orientation differences. A second piece of evidence for a perceptual contribution is that serial dependence is reduced when objects pass behind an occluder and reappear at an inconsistent (as compared with consistent) location (Liberman, Zhang, & Whitney, 2016). This finding again implicates visual continuity in serial dependence, independent of the response decision. Finally, the serial dependence effect is correlated with changes in neuroimaging signals of primary visual cortex (St. John-Saaltink, Kok, Lau and de Lang, 2016), This neural correlate was spatially selective: if the right Gabor patch was probed on the previous trial and the left probed on the current trial there was no correlation between the behavioral or neural data.

What is the evidence for a non-perceptual contribution to the serial dependence phenomenon? A post-perceptual interpretation of serial dependence was proposed by Fritsche et al. (2017). In their Experiment 3, they alternated between adjustment trials (participants saw a Gabor patch followed by a bar that they adjusted to match the stimulus orientation) with same-different judgment trials (participants saw two stimuli and needed to report whether they were the same or different). Fritsche et al. reasoned that if the serial dependence effects are perceptual, they should appear on the same-different judgments, a reporting method that is more bias free than the method of adjustment (Schneider, 2006). However, they found that orientation perception on same-different trials was actually repulsed from the previous stimulus’s orientation, a spatially specific effect that is consistent with previous research (Campbell & Maffei, 1971; Gibson & Radner, 1935). Fritsche et al. (2017) also observed that the magnitude of the serial dependence effect was increased when the delay between perception and judgment was increased (see also: Bliss, Sun, & D’Esposito, 2017). This observation is inconsistent with a mechanism that enhances the spatiotemporal continuity of perception. Altogether, the data of Fritsche et al. demonstrate (i) a spatially specific perceptual effect that is consistent with established perceptual phenomena and (ii) a spatially non-specific post-perceptual bias which resembles the original serial dependence effect.

Our current data are also inconsistent with a purely perceptual account of serial dependence. The serial dependence effect was unaffected when stimuli changed color and/or location across trials. If a continuity field biases perception of distinct objects as if they were the same object, this would appear to be disadvantageous for accurately comparing and successfully distinguishing objects. Additionally, our data indicated that serial dependence was observed even for consecutive stimuli with very different orientations. Altogether, these findings – spatial non-specificity, object non-specificity, and serial dependence for dissimilar features – are most readily accounted for by a model positing an interaction between perceptual features and internal response variables. An example of a toy model that captures the relevant phenomena is described in the Appendix. The model describes the relationship between the probability distributions over stimulus orientation (*s_n_*(*θ*)) and response distributions (*r_n_*(*θ*)) across all trials.

Our simple model would need to be extended to explain why serial dependence is reduced when an object moves behind an occluder and reappears at an inconsistent location (Liberman et al., 2016). We propose that the internal response bias is reset at so-called “event boundaries”, which occur when a surprising event occurs in the environment. This possibility is supported by findings of cognitive effects being eliminated or reversed if the two relevant events are more strongly differentiated in some way (Akyürek, Taffanin, & Hommel, 2008; Pfister, Kiesel, & Mechler, 2010) and is also consistent with the literature on event boundaries in learning and memory (e.g. Botvinick, 2012). One novel prediction of this model is that the serial dependence effect should be larger following runs of large response errors in the same direction as the systematic error would accumulate in the response variable. To test this prediction, an experiment would need to be designed to induce larger and more consistent response errors than observed in the standard serial dependence design.

The simple model we have proposed could also be extended to be an optimal integration model (e.g. see the Bayesian integration model proposed by Cicchini, Anobile, & Burr (2014)). In this case, the parametrization of the combination function *f*(*s*(*θ*), *r*(*θ*)) could be inferred based on the precision of estimates of *s* and *r*. The model could be improved to account for how the representation of stimulus and response variables vary over time following the stimulus offset. Fritsche Mostert, & de Lange (2017) demonstrated that the serial dependence effect increased as the duration between stimulus presentation and the response increased. They interpreted this finding as evidence against a perceptual explanation of the serial dependence effect as the perceptual evidence is the same in both cases and evidence that during the retention interval the working memory bias becomes increasingly biased towards the previous stimuli (Huang & Sekuler, 2010). Such an interpretation is certainly possible within the current model if the contribution of *r*_1_(*θ*) increases relative to the contribution of *s*_1_(*θ*) as the interval between the stimulus and response increases (Pertzov, Bays, Joseph, & Hussain, 2013; Wei, Wang, & Wang, 2012), causing the response threshold to be passed at an orientation further away from the true orientation.

A noteworthy aspect of the current design is that we manipulated perceptual features that belonged exclusively to the figure (as opposed to the ground) of the display. It remains possible that manipulating background features could impact the serial dependence effect: the visual system should not apply a continuity field to two objects appearing within different contexts (e.g. similar looking objects in different rooms). Indeed, background color changes have been shown to reduce or eliminate the serial dependence (Kiyonaga, Manassi, D’Esposito, Whitney, 2017).

Of course, changes in background features (e.g. room features, which indicate the current context for perception and action) could also be expected to reset internal decision/response variables, so the observation of background-feature effects is not diagnostic between perceptual and non-decisional models of the serial dependence effect. Still, the different contributions of figure and ground changes to the serial dependence effects remains an intriguing line of research.

In summary, in three experiments we tested the object specificity of serial dependence. We reasoned that if the serial dependence effect arises from perceptual mechanisms, such as a perceptual continuity field, then it should only transfer between stimuli that could be reasonably inferred to be the same object across time. In contrast to this prediction, we found no evidence of the serial dependence effect being object specific nor spatially specific when objects are differentiated by color, location, or both color and location simultaneously. This non-specificity raises the possibility that the serial dependence effect is not simply perceptual.

We conclude that the serial dependence effect is unlikely to be explained by a perceptual continuity field alone. Instead, it could result from interactions between perceptual features and internal response variables. While there is likely a perceptual contribution to the process producing serial dependence, we note that perceptual systems must balance the need to integrate stimuli with the need to differentiate them: both over-integration and the over-differentiation of incoming visual input are functionally problematic (e.g. Kiyonaga, Scimeca, Bliss, & Whitney, 2017). Also, while Fischer and Whitney (2014) suggested that a continuity field may be the mechanism underlying object file maintenance, our findings cast doubt on this possibility. Object updating studies commonly find that color and location link objects from one moment to the next (Hilchey et al., 2017a, 2017b; Hommel, 1998; Hommel & Colzato, 2004; Hommel, Memelink, Zmigrod, & Colzato, 2013; Quinlan, 1998), but color and location have little effect on the serial dependence thought to arise from the continuity field.

Future research should investigate the boundary conditions under which integration and differentiation of features are applied across objects. One possibility is that the object-oriented ventral stream applies a serial dependence mechanism that helps maintain object continuity while the action-oriented dorsal visual stream applies a mechanism that heightens sensitivity to potentially action relevant stimulus changes (Goodale, 2014; Goodale & Milner, 1992). Such phenomena at the border of perception and action are of special relevance for understanding mental function and behavior in a continuously changing world.

## Appendix

We model the behavioral judgment on each trial of our task as a joint function of perceptual features and an internal response bias (Figure 7). The immediately perceived stimulus features are described by a continuous function, *s_n_*(*θ*), representing the strength of each orientation, *θ*, in the stimulus presented on trial *n*. The internal response bias is likewise a continuous function, *r_n_*(*θ*), describing the internal strength (or response bias) on that trial for each orientation, *θ*. The behavioral judgment, *j_n_*, that is produced on each trial is sampled with probability *p*(*j_n_* = *θ*) = *f*(*s_n_*(*θ*),*r_n_*(*θ*)) where *f* is a probability function that increases for increasing *s* and for increasing *r*. In the simplest case, *f*(*s*(*θ*), *r*(*θ*)) ∝ *s*(*θ*) + *r*(*θ*), with a normalization to ensure that *f* sums to unity. However, this simple averaging would not reproduce the phenomenon that large orientation differences (e.g. 75°) exhibit larger serial dependence than intermediate orientation differences (e.g. 30°). To capture this phenomenon, we can use a function of the form *f*(*s*(*θ*),*r*(*θ*)) ∝ *s*(*θ*)(*ϵ* + *r*(*θ*)), where *ϵ* is typically << 1.

The model makes two key assumptions. First, we assume that when a stimulus of orientation *θ*\* is presented on trial *n*, then *s_n_*(*θ*) becomes a symmetric distribution (such as a Gaussian or central Cauchy distribution) centered on *θ*\*. Second, we assume that on each trial, the internal response bias^1^ is updated as a linear combination of its prior state and the prior stimulus: *r_n_*(*θ*) = *α r*_*n*−1_(*θ*) + (1 – *α*)*s*_*n*−1_(*θ*). Here, *α* is a parameter between 0 and 1 that modulates the timescale of serial dependence. When *α* = 0, then the response bias on each trial will be modulated only by the stimulus on the immediately preceding trial; when *α* > 0, then serial dependence may be observed over many consecutive trials.

Figure 8 illustrates how this model works. On the first trial the internal response bias, *r*_1_(*θ*) is a uniform function. Upon presentation of a stimulus with true orientation 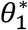, the stimulus activation, *s*_1_(*θ*), becomes a Gaussian distribution centered on 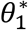. The function describing the behavioral response *p*_1_(*θ*) is then also approximately Gaussian and centered on 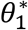 (panel a), because the response bias function, *r*_1_(*θ*), is flat. However, for the trial following, the response bias function, *r*_2_(*θ*), is no longer flat, but is instead centered on 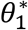 (panel b). Therefore, if the stimulus orientation on the second trial, 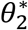, is greater than (clockwise from) the stimulus on the first trial, then the behavioral response function *p*_2_(*θ*) will reflect a combination of the previous stimulus and current stimulus, and will be centered on a value between 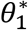 and 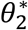. This shifting of the mode and expectation of *p*_2_(*θ*) manifests as a serial dependence effect (panel c). Note that the serial dependence effect decreases in magnitude when consecutive stimuli are very different from each other, because of the functional form assumed for *f*(*s*(*θ*), *r*(*θ*)).

**Figure A1.**
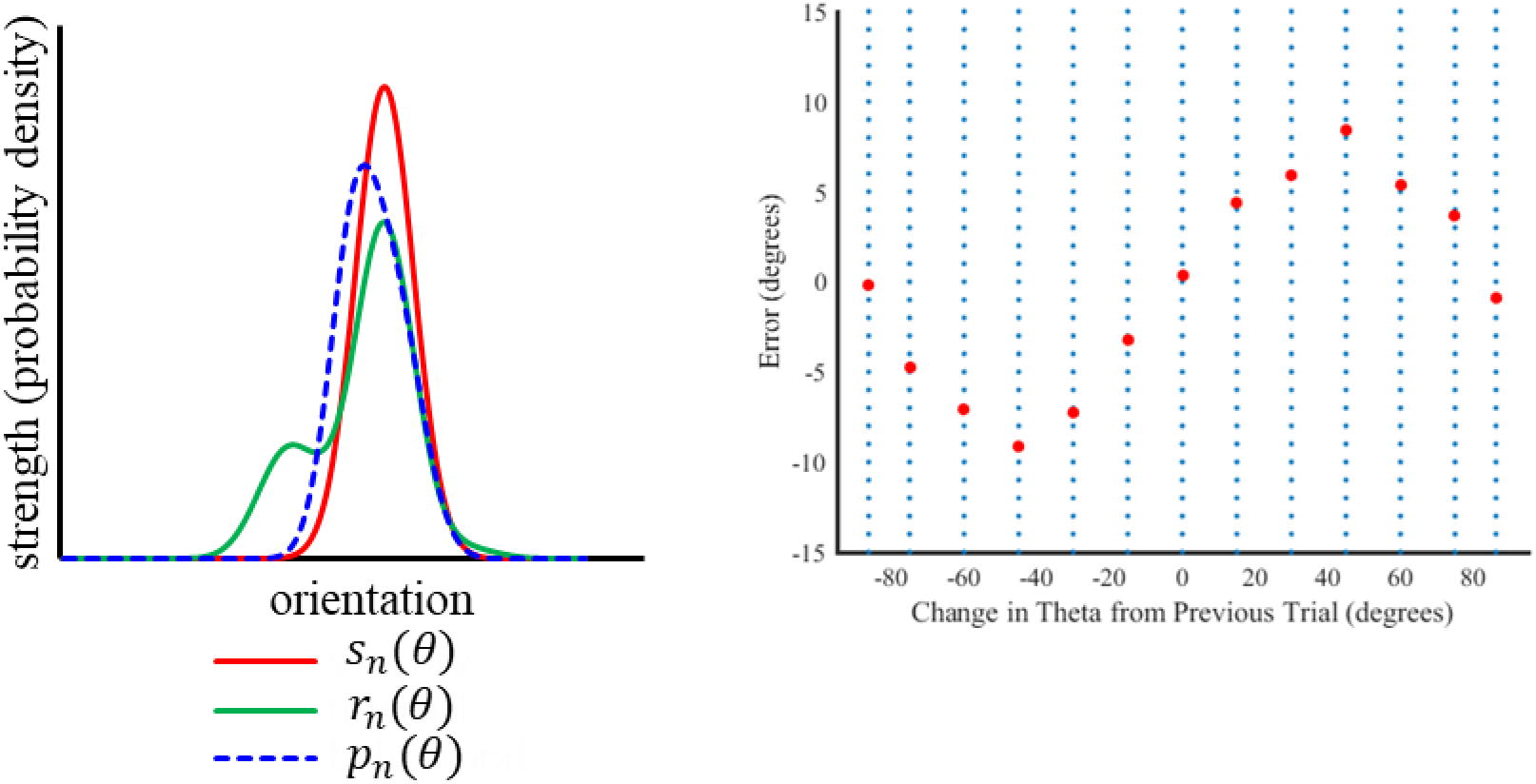
A model of the serial dependence effect in which serial dependence arises from interactions between stimulus-driven sensory response and abiasn internal response which is influenced by the previous trial’s response.

1 A critical component of this model is that we assume that the response selection process, and not executing the response, that leads to the bias. This assumption is based on a substantial amount of theory and research regarding the interaction of stimulus and response factors (Gozli, Huffman, & Pratt, 2016; Hommel, Müsseler, Aschersleben, & Prinz, 2001; Kunde, 2001; Schumacher & Hazeltine, 2016).

## References

Akyürek, E. G., Toffanin, P., & Hommel, B. (2008). Adaptive control of event integration. Journal of Experimental Psychology: Human Perception and Performance, 34, 569–577.

Botvinick, M. M. (2012). Hierarchical reinforcement learning and decision making. Current Opinion in Neurobiology, 22, 956–962.

Campbell, F. W., & Maffei, L. (1971). The tilt after-effect: A fresh look. Vision Research, 11, 833–840.

Carmel, T., & Lamy, D. (2014). The same-location cost is unrelated to attentional settings: An object-updating account. Journal of Experimental Psychology: Human Perception and Performance, 40(4), 1465.

Chen, Z. (2009). Not all features are created equal: Processing asymmetries between location and object features. Vision Research, 49(11), 1481–1491.

Cicchini, G. M., Anobile, G., & Burr, D. C. (2014). Compressive mapping of number to space reflects dynamic encoding mechanisms, not static logarithmic transform. Proceedings of the National Academy of Sciences of the United States of America, 111, 7867–7872.

Clark, A. (2013). Whatever next? Predictive brains, situated agents, and the future of cognitive science. The Behavioral and Brain Sciences, 36, 181–204.

Fischer, J., & Whitney, D. (2014). Serial dependence in visual perception. Nature Neuroscience, 17, 738–743.

Friston, K., & Kiebel, S. (2009). Predictive coding under the free-energy principle. Philosophical Transactions of the Royal Society of London B: Biological Sciences, 364, 1211–1221.

Fritsche, M., Mostert, P., & de Lange, F. P. (2017). Opposite effects of recent history on perception and decision. Current Biology, 27, 590–595.

Folk, C. L., Remington, R. W., & Johnston, J. C. (1992). Involuntary covert orienting is contingent on attentional control settings. Journal of Experimental Psychology: Human perception and performance, 18(4), 1030.

Gibson, J. J., & Radner, M. (1937). Adaptation, after-effect and contrast in the perception of tilted lines. I. Quantitative studies. Journal of Experimental Psychology, 20, 453–467.

Golomb, J. D., Kupitz, C. N., & Thiemann, C. T. (2014). The influence of object location on identity: A “spatial congruency bias”. Journal of Experimental Psychology: General, 143(6), 2262.

Goodale, M.A. (2014) How (and why) the visual control of action differs from visual perception. Proceedings of the Royal Academy of Science: B, 281(20140337). doi:10.1098/rspb.2014.0337

Goodale, M. A., & Milner, A. D. (1992). Separate visual pathways for perception and action. Trends in Neurosciences, 15, 20–25.

Gozli, D. G., Huffman, G., & Pratt, J. (2016). Acting and anticipating: Impact of outcome-compatible distractor depends on response selection efficiency. Journal of Experimental Psychology: Human Perception and Performance, 42, 1601–1614.

Hilchey, M. D., Rajsic, J., Huffman, G., & Pratt, J. (2017a). Intervening response events between identification targets do not always turn repetition benefits into repetition costs. Attention, Perception, & Psychophysics, 79(3), 807–819.

Hilchey, M. D., Rajsic, J., Huffman, G., & Pratt, J. (2017b). Response-mediated spatial priming despite perfectly valid target location cues and intervening response events. Visual Cognition, 1–15.

Hilchey, M. D., Rajsic, J., Huffman, G., Klein, R. M., & Pratt, J. (2018). Dissociating Orienting Biases From Integration Effects With Eye Movements. Psychological science, 0956797617734021.

Hommel, B. (1998). Event files: Evidence for automatic integration of stimulus-response episodes. Visual Cognition, 5(1–2), 183–216.

Hommel, B. (2002). Responding to object files: Automatic integration of spatial information revealed by stimulus-response compatibility effects. The Quarterly Journal of Experimental Psychology Section A, 55(2), 567–580.

Hommel, B. (2005). How much attention does an event file need?. Journal of Experimental Psychology: Human Perception and Performance, 31(5), 1067.

Hommel, B., & Colzato, L. (2004). Visual attention and the temporal dynamics of feature integration. Visual Cognition, 11(4), 483–521.

Hommel, B., Müsseler, J., Aschersleben, G., & Prinz, W. (2001). The Theory of Event Coding (TEC): A framework for perception and action planning. The Behavioral and Brain Sciences, 24, 849–937.

Huang, J., & Sekuler, R. (2010). Attention protects the fidelity of visual memory: Behavioral and electrophysiological evidence. Journal of Neuroscience, 30, 13461–13471.

Huffman, G., Antinucci, V. M., & Pratt, J. (2018). The illusion of control: Sequential dependencies underlie contingent attentional capture. Psychonomic bulletin & review, 1–7.

JASP Team (2018). JASP (Version 0.9)[Computer software].

John-Saaltink, E. S., Kok, P., Lau, H. C., & De Lange, F. P. (2016). Serial dependence in perceptual decisions is reflected in activity patterns in primary visual cortex. Journal of Neuroscience, 36(23), 6186–6192.

Kahneman, D., Treisman, A., & Gibbs, B. J. (1992). The reviewing of object files: Object-specific integration of information. Cognitive Psychology, 24, 175–219.

Keizer, A. W., Hommel, B., & Lamme, V. A. (2015). Consciousness is not necessary for visual feature binding. Psychonomic bulletin & review, 22(2), 453–460.

Kiyonaga, A., Manassi, M., D’Esposito, M., & Whitney, D. (2017). Context transitions modulate perceptual serial dependence. Journal of Vision, 17(10), 92–92.

Kiyonaga, A., Scimeca, J. M., Bliss, D. P., & Whitney, D. (2017). Serial Dependence across perception, attention, and memory. Trends in Cognitive Sciences, 21, 493–497.

Kleiner, M., Brainard, D., Pelli, D., Ingling, A., Murray, R., & Broussard, C. (2007). What’s new in Psychtoolbox-3. Perception, 36, ECVP Abstract Supplement.

Kunde, W. (2001). Response-effect compatibility in manual choice reaction tasks. Journal of Experimental Psychology: Human Perception and Performance, 27, 387–394

Liberman, A., Fischer, J., & Whitney, D. (2014). Serial dependence in the perception of faces. Current Biology, 24, 2569–2574.

Liberman, A., Zhang, K., & Whitney, D. (2016). Serial dependence promotes object stability during occlusion. Journal of Vision, 16, 1–10.

Pfister, R., Kiesel, A., & Melcher, T. (2010). Adaptive control of ideomotor effect anticipations. Acta Psychologica, 135, 316–322.

Pertzov, Y., Bays, P. M., Joseph, S., & Husain, M. (2013). Rapid forgetting prevented by retrospective attention cues. Journal of Experimental Psychology: Human Perception and Performance, 39, 1224–1231.

Rogers, T. T., & McClelland, J. L. (2014). Parallel distributed processing at 25: Further explorations in the microstructure of cognition. Cognitive science, 38, 1024–1077.

Schneider, K. A. (2006). Does attention alter appearance?. Perception & Psychophysics, 68, 800–814.

Schoeberl, T., Ditye, T., & Ansorge, U. (2018). Same-location costs in peripheral cueing: The role of cue awareness and feature changes. Journal of Experimental Psychology: Human Perception and Performance, 44(3), 433.

Schumacher, E. H., & Hazeltine, E. (2016). Hierarchical task representation: Task files and response selection. Current Directions in Psychological Science, 25, 449–454.

Spadaro, A., He, C., & Milliken, B. (2012). Response to an intervening event reverses nonspatial repetition effects in 2AFC tasks: Nonspatial IOR?. Attention, Perception, & Psychophysics, 74, 331–349.

Taubert, J., & Alais, D. (2016). Serial dependence in face attractiveness judgements tolerates rotations around the yaw axis but not the roll axis. Visual Cognition, 24, 103–114.

Taubert, J., Van der Burg, E., & Alais, D. (2016). Love at second sight: Sequential dependence of facial attractiveness in an on-line dating paradigm. Scientific reports, 6, 1–5.

Theeuwes, J. (1992). Perceptual selectivity for color and form. Attention, Perception, & Psychophysics, 51, 599–606.

Treisman, A. M., & Gelade, G. (1980). A feature-integration theory of attention. Cognitive Psychology, 12, 97–136.

van Dam, W. O., & Hommel, B. (2010). How object-specific are object files? Evidence for integration by location. Journal of Experimental Psychology: Human Perception and Performance, 36(5), 1184.

Wei, Z., Wang, X. J., & Wang, D. H. (2012). From distributed resources to limited slots in multiple-item working memory: a spiking network model with normalization. Journal of Neuroscience, 32, 11228–11240.

Xia, Y., Leib, A. Y., & Whitney, D. (2016). Serial dependence in the perception of attractiveness. Journal of Vision, 16, 1–8.

